# Next-generation sequencing-based liquid biopsy can be used for detection of residual disease and cancer recurrence monitoring in dogs

**DOI:** 10.1101/2023.09.08.556935

**Authors:** Angela L. McCleary-Wheeler, Patrick C. Fiaux, Carlos A. Ruiz-Perez, Lisa M. McLennan, John A. Tynan, Susan C. Hicks, Jill M. Rafalko, Daniel S. Grosu, Jason Chibuk, Allison L. O’Kell, Todd A. Cohen, Brian K. Flesner, Ilya Chorny, Dana W.Y. Tsui, Kristina M. Kruglyak, Andi Flory

**Affiliations:** PetDx, Research Programs, La Jolla, CA; PetDx, Information Technology, La Jolla, CA; PetDx, Analytical Production, La Jolla, CA; PetDx, Medical & Clinical Affairs, La Jolla, CA

## Abstract

**Objective:** The purpose of this study was to evaluate the performance of a next-generation sequencing-based liquid biopsy test for cancer monitoring in dogs.

**Samples:** Pre- and post-operative blood samples were collected prospectively from dogs with confirmed cancer diagnoses originally enrolled in the CANcer Detection in Dogs (CANDiD) study. A subset of these dogs also had longitudinal blood samples collected for recurrence monitoring.

**Methods:** All patients had a pre-operative blood sample collected (after diagnosis but prior to surgical intervention) in which a cancer signal was detected, and had at least one post-operative sample collected. Clinical data were collected for all patients and used to assign a clinical disease status for each follow-up visit.

**Results:** Following excisional surgery, in the absence of clinical residual disease at the post-operative visit, patients with *Cancer Signal Detected* results at that visit were 1.95-times as likely to have clinical recurrence within 6 months compared to patients with *Cancer Signal Not Detected* results. In the subset of patients with longitudinal liquid biopsy samples that had clinical recurrence documented during the study period, 73% (8/11; 95% CI: 39 – 93%) of patients had *Cancer Signal Detected* in blood prior to or concomitant with clinical recurrence; in the 6 patients where molecular recurrence was detected prior to clinical recurrence, the median lead time was 168 days (range: 47 – 238).

**Clinical Relevance:** Next-generation sequencing-based liquid biopsy is a non-invasive tool for cancer monitoring in dogs that can be used as an adjunct to current standard-of-care clinical assessment methods.

## Introduction

After a patient is diagnosed with cancer and receives therapy, the current standard of care relies heavily on physical examination and imaging to assess disease status and monitor for cancer recurrence (local recurrence of cancer and/or regional/distant metastasis; hereafter referred to as cancer recurrence). However, these monitoring tools may not be sensitive enough to detect the presence of residual disease following excisional surgery, or early evidence of cancer recurrence following therapeutic intervention. Additionally, access to certain imaging modalities may be limited in some care settings, and evaluation by imaging may require sedation or anesthesia, posing risks to the patient.

In human oncology, next-generation sequencing-based (NGS-based) liquid biopsy tests have been incorporated into clinical practice for cancer monitoring in the post-diagnosis setting. The detection of cancer-associated genomic alterations (cancer signal) in a patient’s blood after treatment is considered a prognostic marker for a variety of human cancer types (solid as well as hematological), including colorectal cancer^1,2^, lung cancer^3^, melanoma^4,5^, bladder cancer^6^, and lymphoma^7^, among others^8–11^. NGS-based liquid biopsy has also been shown to detect cancer signal prior to clinical recurrence in human patients, ^1,11^ however, such evidence has not yet been established in veterinary medicine.

The following analyses evaluated the performance of a blood-based liquid biopsy test that has been clinically validated for cancer detection in dogs^12^, as an adjunct tool for cancer monitoring. The study hypothesis was that, in dogs who underwent excisional surgery, patients with a *Cancer Signal Detected* (CSD) result at a post-operative visit (in the presence of no clinical residual disease) would have a higher risk of clinical recurrence within 6 months compared to those with a *Cancer Signal Not Detected* (CSND) result. A secondary objective was to retrospectively determine whether liquid biopsy could detect molecular recurrence of disease prior to or concomitant with clinical recurrence in a subset of patients that were longitudinally monitored.

## Materials and Methods

### Assessment of liquid biopsy for the detection of residual disease

The study population comprised 52 dogs originally enrolled in Protocol 301 of the CANcer Detection in Dogs (CANDiD) study **(Figure 1)**^12,13^. The CANDiD study was a clinical validation study of NGS-based liquid biopsy in dogs and enrolled presumably cancer-free and cancer-diagnosed dogs into three protocols (101, 201, and 301)^12^. Protocol 301 included dogs scheduled for biopsy or surgery due to suspected or known cancer. For the current study, each patient had a confirmed diagnosis of at least one primary solid tumor malignancy and underwent excisional surgery of their cancer along with standard-of-care adjuvant therapy (which may have included chemotherapy, radiation therapy, immunotherapy, etc.) at their managing veterinarian’s office. Each patient had a baseline (pre-operative) blood sample drawn for NGS-based liquid biopsy testing immediately prior to surgical excision of their cancer; all dogs had CSD results at baseline. Patients with CSD results at baseline were chosen because cancer detection by liquid biopsy is known to vary depending on the cancer type present in the patient^12^; therefore, the detection of a cancer signal at baseline (prior to any therapeutic intervention) in a cancer-diagnosed patient suggests that the test can be subsequently used in that patient as a tool for cancer monitoring. Baseline samples were submitted between November 2019 and March 2022, with additional samples collected at follow-up visits as described below. Samples were stored for liquid biopsy testing as previously described.^12^

**Figure 1:**
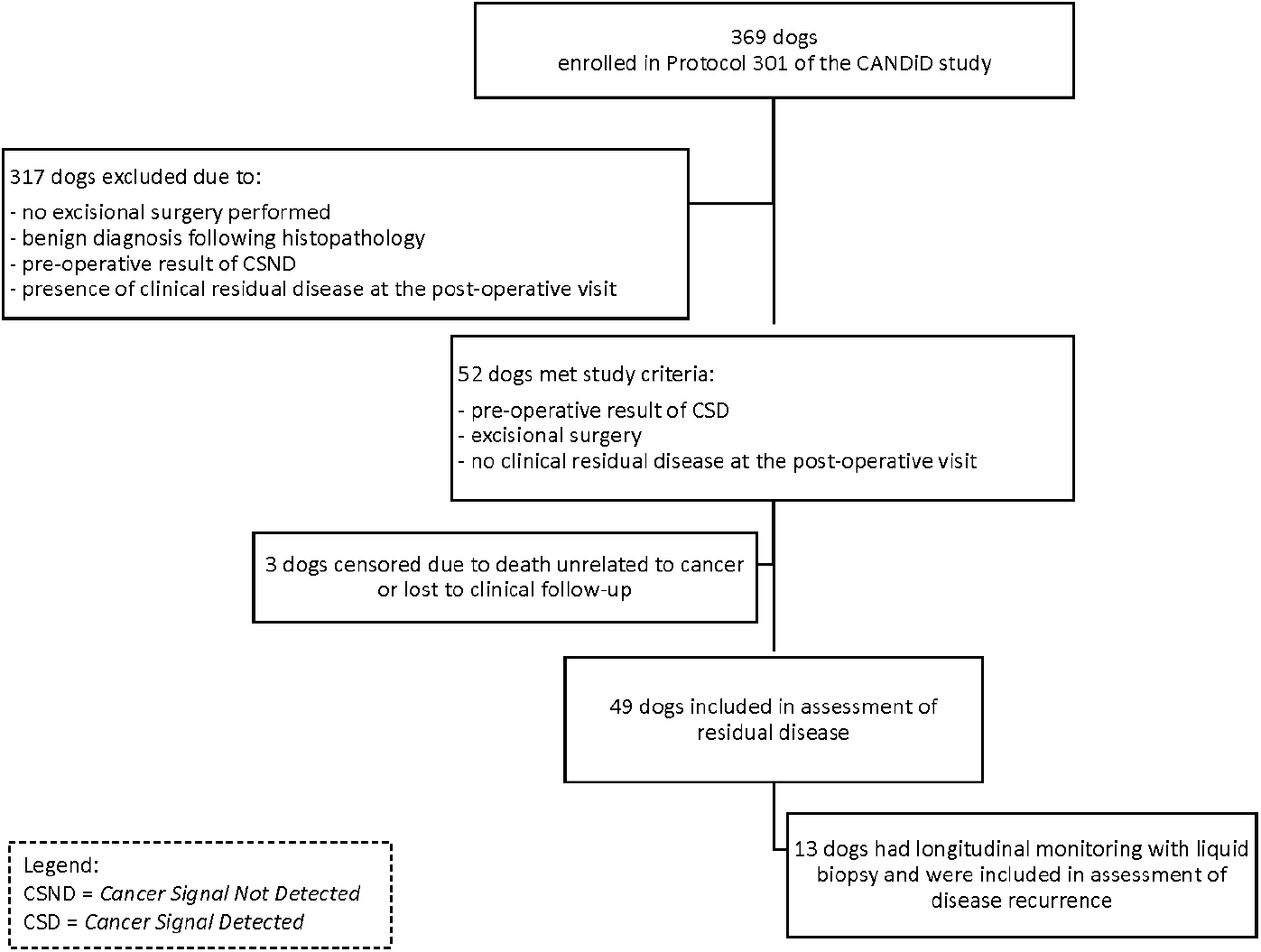
Disposition of patients from enrollment in Protocol 301 of the CANcer Detection in Dogs (CANDiD) study through the current study which evaluated the performance of a next-generation sequencing-based liquid biopsy test as an adjunct tool for cancer monitoring in dogs – specifically for the detection of residual disease following surgery and detection of cancer recurrence.

In addition to the baseline blood sample, each patient had a follow-up blood sample collected after surgery at the post-operative visit (3-34 days following surgery), and had no clinical residual disease documented at that visit **(Figure 2)**. Histopathology reports were reviewed for margin measurement and completeness of excision (when available) and cases were classified as complete or incomplete based on pathologist report; margins were further categorized as complete and wide if histologically measured ≥5 mm and complete but narrow if histologically measured <5 mm. Excision was considered complete for patients having amputation for a tumor of the long bone or complete organ removal for a tumor confined to an organ (i.e, splenectomy for splenic tumor or nephrectomy for renal tumor) unless specifically denoted as incomplete on the pathology report. Patients that had more than one cancer type were categorized as complete if both cancers were noted to be completely excised, and incomplete if either tumor was noted to be incompletely excised. Beyond the post-operative visit, additional clinical outcome data were retrospectively collected for each patient in this cohort through a final patient outcome report completed by the treating veterinarian. Information collected included: whether a patient had developed clinical evidence of recurrence; the documented date of recurrence; if the patient had died; and the date and cause of death.

**Figure 2:**
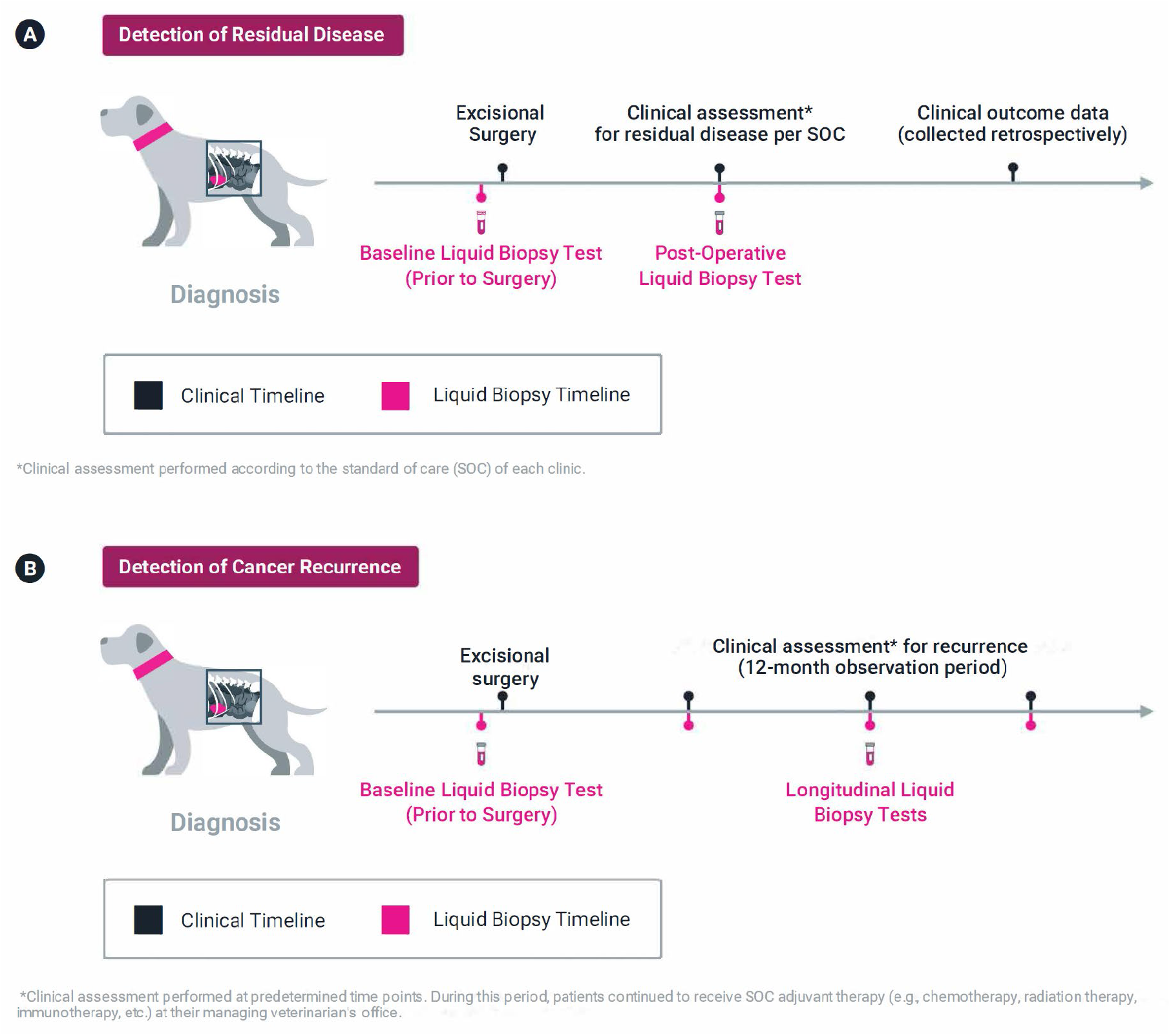
Timeline of clinical evaluations and blood sample collections for the cancer-diagnosed patients from which detection of residual disease and detection of disease recurrence was assessed. (A) 52 patients met criteria to assess the test’s ability to detect residual disease following excisional surgery, and (B) a subset of 13 patients met criteria to assess the test’s ability to detect cancer recurrence.

The 52 patients were stratified by their liquid biopsy test results at the post-operative visit into two groups, CSD and CSND, and were compared for their (i) rate of recurrence and (ii) recurrence-free survival (RFS), in both cases within 6 months of excisional surgery. RFS results were expressed using Kaplan-Meier (KM) curves. Patients who were lost to follow-up or died due to non-cancer related causes within 6 months (n=3) were censored in all calculations.

### Assessment of liquid biopsy for the detection of cancer recurrence

Of the aforementioned 52 patients that had surgical resection of their cancer, a subset of 13 patients had additional blood samples collected longitudinally during follow-up visits on a pre-defined schedule at 1, 3, 6, 9, and 12 months following the baseline visit **(Figure 2)**. Not all patients completed all predefined visits and, in some cases, additional samples were obtained at non-predefined visits, e.g., when the treating veterinarian noted a change in a patient’s clinical disease status. At each follow-up visit, the patient’s clinical disease status was evaluated by a specialist (in surgery or oncology) using available standard-of-care methods such as physical exam, bloodwork, and routine imaging (radiography and/or ultrasound); and the treating veterinarian assigned a standardized clinical disease status using cRECIST criteria for solid tumors^14^, as follows: progressive disease (PD), stable disease (SD), partial response (PR), and complete response (CR); a no evidence of disease (NED) status designation was considered synonymous with CR. These designations, and supporting clinical records, were subsequently reviewed by an ACVIM board-certified veterinary medical oncologist at the laboratory (PetDx, La Jolla, CA) to confirm that the treating veterinarian’s clinical disease status designation was aligned with the standardized criteria at each visit. The clinical disease status assigned at each visit was further used to categorize each visit as either presence of clinical disease (corresponding to SD, PR, or PD) or absence of clinical disease (corresponding to CR). The clinical team at PetDx that reviewed the clinical disease status designations were blinded to the liquid biopsy test results.

### Statistical analysis

Normality of data was assessed for age and weight of study patients, and for molecular recurrence lead-times using Shapiro-Wilk tests and visual inspection using QQ-plots. Comparison of 6-month cancer recurrence in patients with CSD versus CSND results was performed using Fisher’s Exact Test. Confidence intervals of proportions were calculated using the Wilson procedure. Recurrence free survival results were expressed using Kaplan-Meier (KM) curves with a hazard ratio calculated by dividing the recurrence rate in patients with CSD results by the recurrence rate in patients with CSND results. Calculations and plotting were performed in R (version 4.3.0) and Python v3.9. A p-value of < 0.05 was considered statistically significant.

## Results

The 52 patients in the study ranged in age from 1.9 to 14.5 years (median 10 years); weights ranged from 9.5 to 60 kg (mean 31.8 kg); 56% of patients were male and 44% were female; 96% of males were neutered and 100% of females were spayed; 42% were purebred and 58% were mixed-breed; 12 cancer types were represented among the patients (10 as solitary diagnoses and 2 as part of multiple cancer diagnoses).

### NGS-based liquid biopsy for detection of residual disease following excisional surgery

Of the 52 unique patients who underwent excisional surgery and had no clinical residual disease at the post-operative visit, three patients were censored due to death unrelated to cancer or because they were lost to clinical follow-up. In the remaining 49 patients **(Table 1)**, those with a CSD result at the post-operative visit were found to have an approximately twofold higher likelihood of recurrence within 6 months (12/15, 80%) compared to patients with a CSND result at that visit (14/34, 41%) (**Figure 3**; *p* = 0.013). Patients with a CSD result at the post-operative visit also had shorter recurrence-free survival within the 6-month observation period compared to those with a CSND result (*p* = 0.015; Hazard Ratio 2.523, 95% CI: 1.161 – 5.484).

**Table 1:**
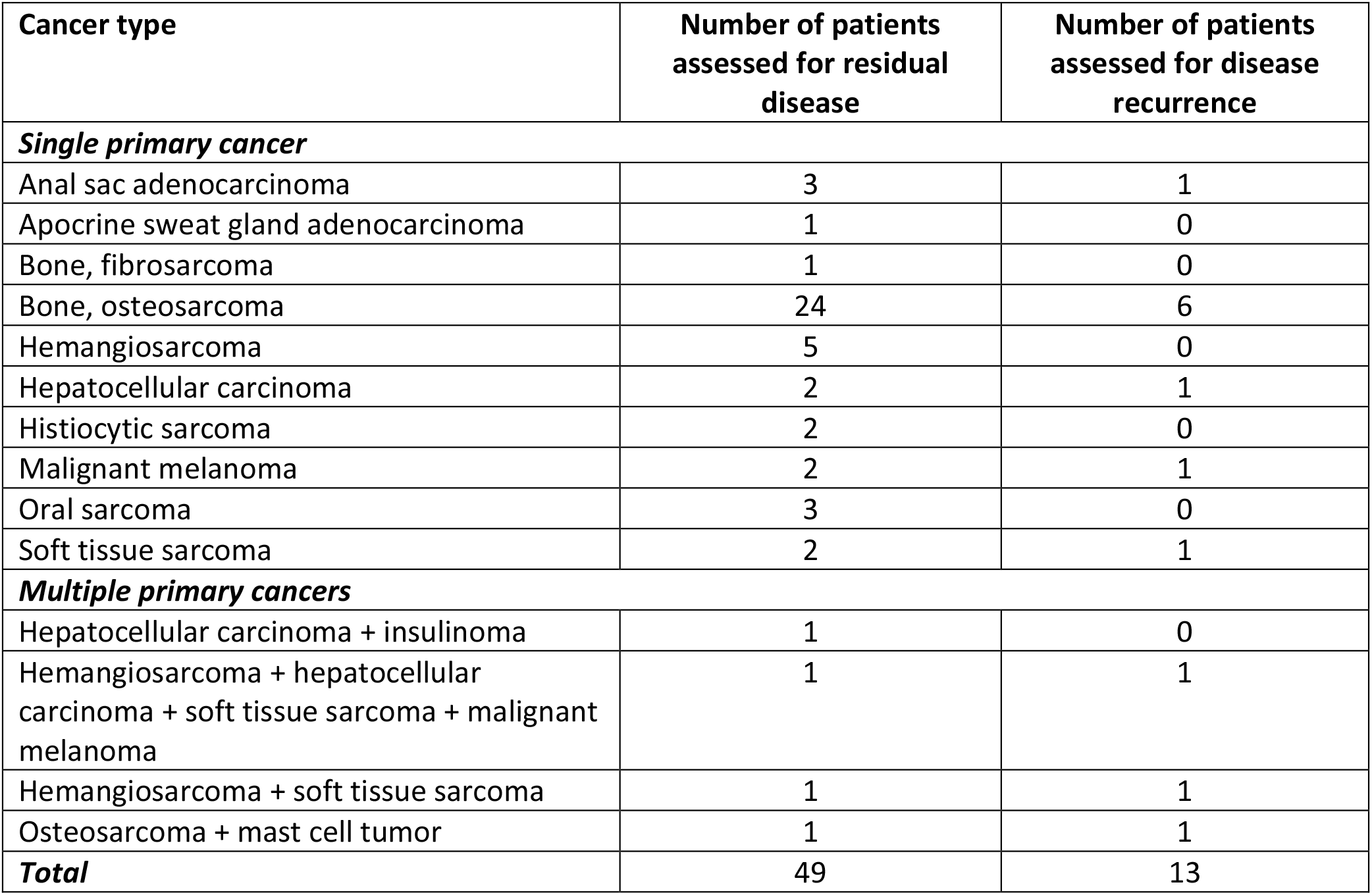
Number of patients assessed for residual disease and disease recurrence by cancer type using NGS-based liquid biopsy.

**Figure 3:**
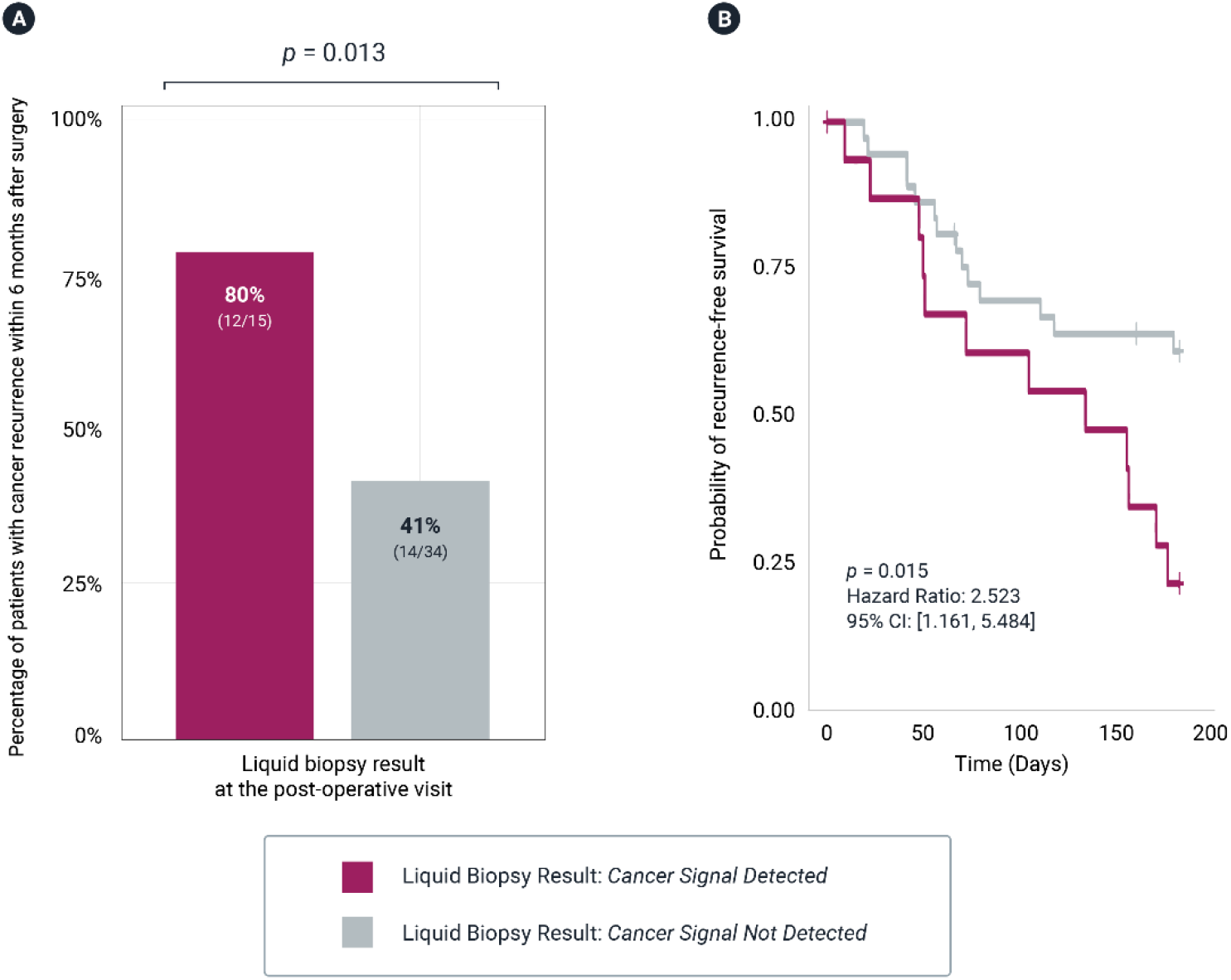
Analysis of outcomes for the 52 unique patients (described in Figure 1) from which detection of residual disease was assessed. Note: Three patients were excluded from final analysis due to death unrelated to cancer or because they were lost to follow-up. (A) Evaluation of the 6-month recurrence rate of patients with post-operative Cancer Signal Detected versus Cancer Signal Not Detected liquid biopsy results. (B) Evaluation of recurrence-free survival over a 6-month period for patients with post-operative Cancer Signal Detected versus Cancer Signal Not Detected liquid biopsy results.

All 49 patients had assessable margins on histopathology reports: 40 had complete margins (including 3 cases with complete and narrow margins <5 cm, 6 with complete and wide margins ≥5 cm, 28 involving amputation or complete organ removal, and 3 with notation of complete margins but no measurements provided) and 9 had incomplete margins (including 1 case of amputation due to intramuscular hemangiosarcoma, but the presence of neoplastic cells continuing to the tissue margins). Based on margin assessment alone, there was no significant difference in cancer recurrence within 6 months, with 50% (20/40) of patients with complete margins and 67% (6/9) of patients with incomplete margins showing recurrence within 6 months (*p* = 0.472). Focusing on the 40 cases with complete margins (which included 25 patients treated with amputation, 20 of which were patients with osteosarcoma), a CSD result at the post-operative visit was associated with a significantly higher rate of cancer recurrence within 6 months (83%; 10/12) compared to cases with complete margins and a CSND result post-operatively (36%; 10/28; *p* = 0.006). When osteosarcoma cases involving amputation were excluded from analysis, 15 cases with complete margins remained, and a post-operative CSD result was still associated with a significantly higher rate of cancer recurrence within 6 months (80%; 4/5) compared to cases with complete margins and CSND results (10%; 1/10; *p* = 0.017).

### NGS-based liquid biopsy for detection of cancer recurrence following therapy

Thirteen of the 52 surgical excision patients had longitudinal monitoring with liquid biopsy, 7 of these dogs had osteosarcoma with amputation (including one dog that had concurrent mast cell tumor involving the same limb). Two patients were reported to be in complete remission throughout the monitoring period (12-months and 9-months, respectively); these dogs had CSND (negative) liquid biopsy results at all follow-up visits. An additional two patients assessed to be in CR (with negative liquid biopsy results) following surgery were noted to have clinical recurrence (progressive disease) at their day 83 and day 84 follow-up visits, respectively. In these two patients, liquid biopsy identified reemergence of cancer signal concomitant with clinical recurrence. The final three patients experienced clinical recurrence at post-op days 79, 277, and 351, but liquid biopsy identified molecular recurrence (i.e., a cancer signal in blood prior to clinical recurrence) in all three cases, with lead times of 47 days, 69 days, and 168 days, respectively **(Figure 4)**.

**Figure 4:**
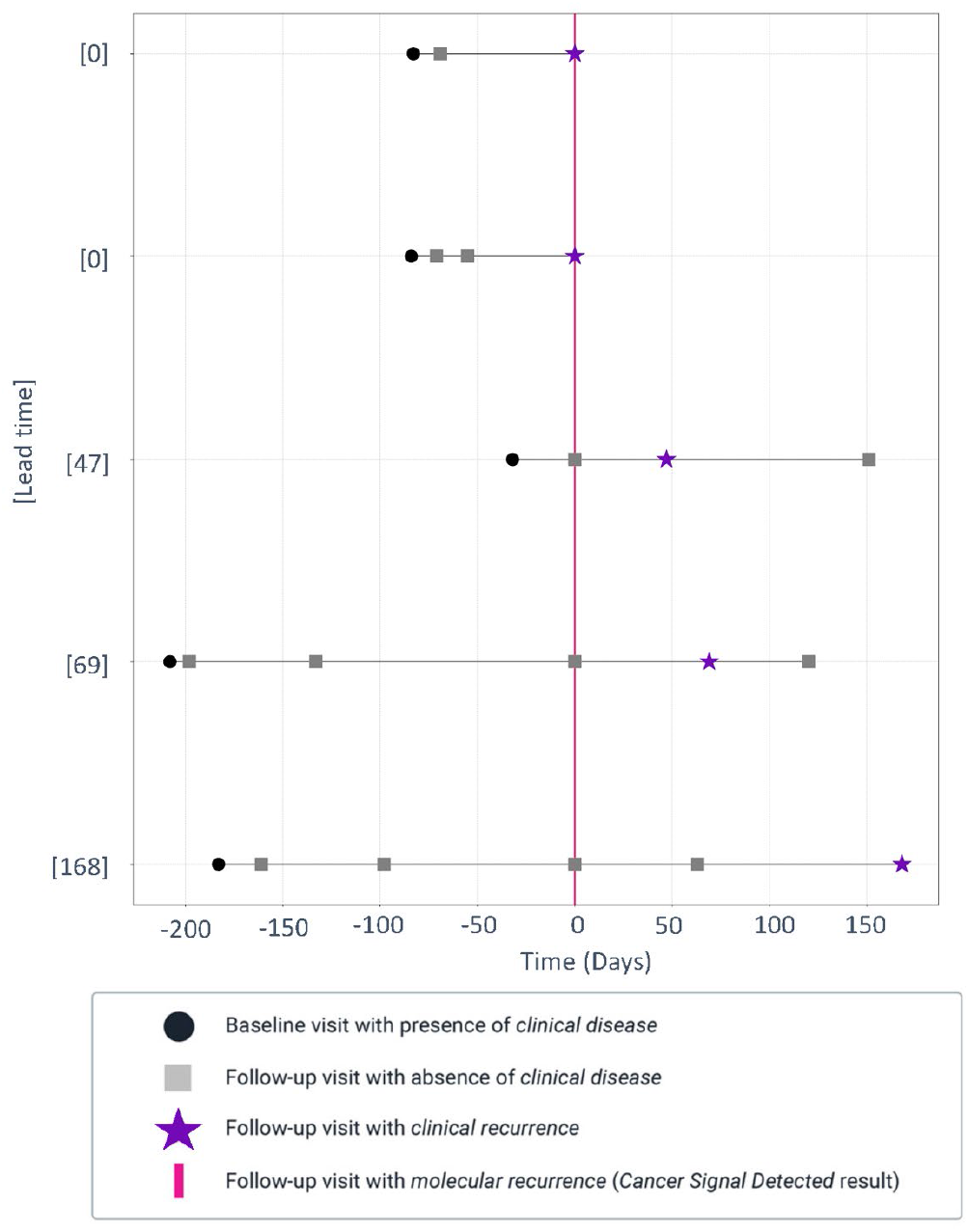
Patient-specific timelines for the seven dogs with osteosarcoma that were longitudinally monitored with liquid biopsy. Two patients were reported to be in complete remission throughout the monitoring period, with Cancer Signal Not Detected liquid biopsy results at all follow-up visits. The other five patients had clinical recurrence during the monitoring period and liquid biopsy detected a cancer signal in blood concomitant with (n=2) or prior to (n=3) clinical recurrence. In cases where liquid biopsy detected molecular recurrence prior to clinical recurrence, the lead times were 47, 69, and 168 days.

Six additional dogs with cancer types other than osteosarcoma had longitudinal monitoring with liquid biopsy. Five of these dogs had documented clinical recurrence (30 to 290 days following surgery). In three of these patients, molecular recurrence was identified via liquid biopsy prior to clinical recurrence (with lead times of 168, 174, and 238 days, respectively; **Table 2**).

**Table 2:** Overview of patients with non-osteosarcoma cancer types that had longitudinal monitoring with NGS-based liquid biopsy. Thee patients had molecular recurrence detected prior to clinical recurrence.

| Cancer type(s) | Number of days from surgery to clinical recurrence | Was molecular recurrence detected? | Lead time (# days from molecular recurrence to clinical recurrence) |
| --- | --- | --- | --- |
| Soft tissue sarcoma | 275 | Yes | 238 |
| Hepatocellular carcinoma | 198 | Yes | 174 |
| Hemangiosarcoma + hepatocellular carcinoma + soft tissue sarcoma + malignant melanoma | 184 | Yes | 168 |
| Anal gland adenocarcinoma | 290 | No | N/A |
| Malignant melanoma | 84 | No | N/A |
| Hemangiosarcoma + soft tissue sarcoma | 30 | No | N/A |

Overall, of the 13 patients that were longitudinally monitored with liquid biopsy, 11 had clinical recurrence of disease documented within the 12-month observation period; 73% (8/11; 95% CI: 39 – 93%) of patients had a CSD in blood concomitant with (n=2) or prior to (n=6) clinical recurrence. In the cases where molecular recurrence was detected ahead of clinical recurrence, the average lead time was 168 days (range: 47 – 238 days).

A case example demonstrating the detection of molecular recurrence by liquid biopsy prior to clinical recurrence involved a 10-year-old spayed female mixed-breed dog diagnosed with osteosarcoma of the right proximal humerus (**Figure 5**). This dog, as all others in this study, had a pre-operative CSD result. After amputation, genomic testing of the tumor tissue revealed genomic alterations concordant with those observed in the pre-operative blood sample. At the post-operative recheck visit a follow-up liquid biopsy test revealed a CSND result. The dog received adjuvant chemotherapy with four doses of carboplatin administered every 3 weeks. At 3-months post-amputation, NED was noted by the treating veterinary oncologist based on physical examination and thoracic radiographs, and simultaneous liquid biopsy continued to show a CSND result. Clinical disease evaluation at 6-months post-amputation (including thoracic radiographs) again documented NED; however, liquid biopsy performed at the same time yielded a CSD result, indicative of molecular recurrence. The CSD result persisted at the 9-month post-surgery evaluation, and at this point, clinical recurrence (PD) was noted with a new, expansile mass on the right 11th rib identified on thoracic radiographs. The patient was rechecked again at 11-months post-surgery and found to have further PD with pulmonary metastasis, and persistence of the CSD liquid biopsy result.

**Figure 5:**
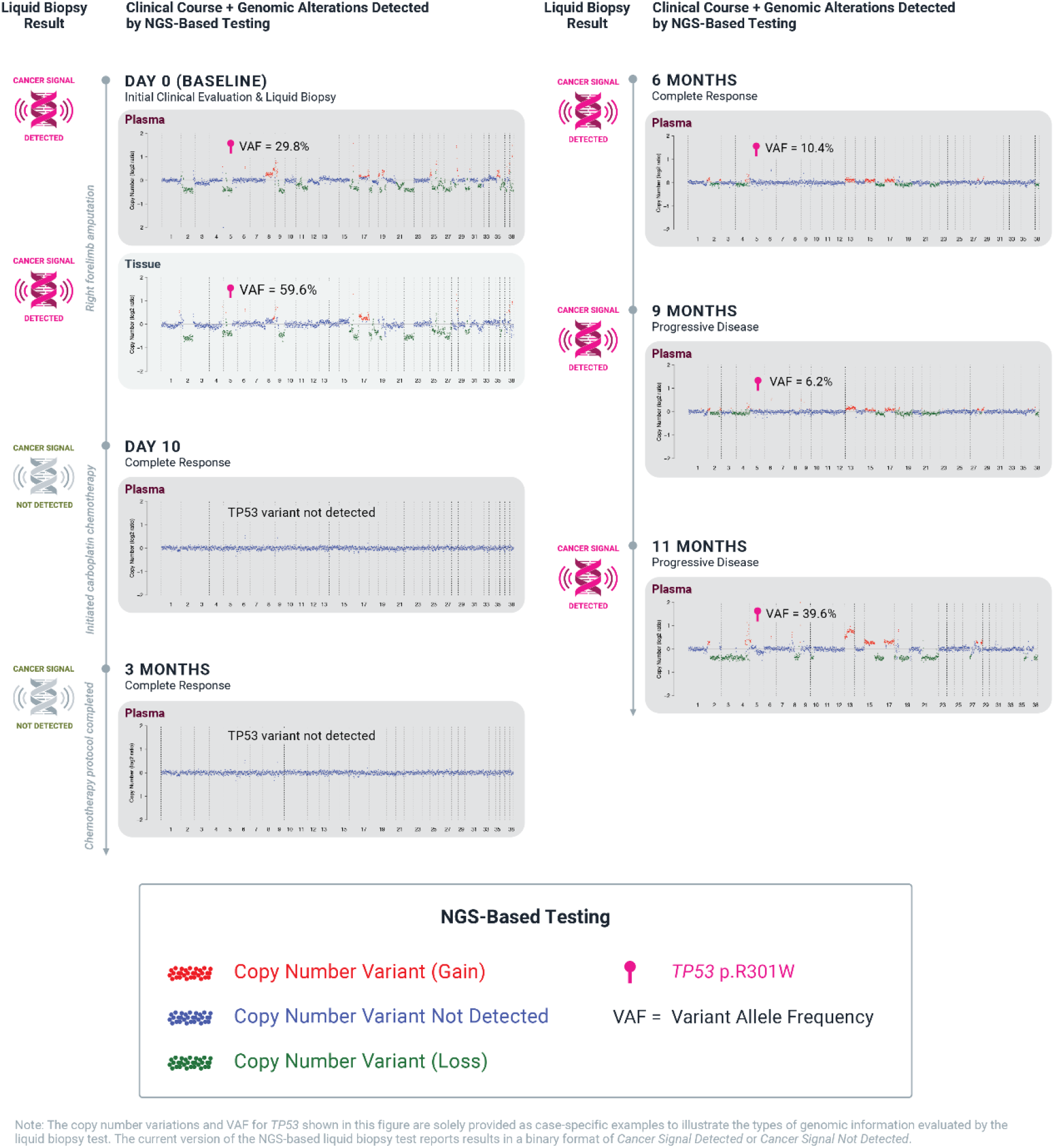
Genomic testing performed in a 10-year-old female spayed mixed-breed dog with a diagnosis of right humeral osteosarcoma. The clinical disease status designation at each visit was based on standard-of-care assessment. The liquid biopsy results are shown as Cancer Signal Detected or Cancer Signal Not Detected in the pre-operative plasma, resected tumor tissue, and plasma collected after the surgery and longitudinally every 2-3 months thereafter.

## Discussion

The results presented herein demonstrate for the first time in veterinary medicine in a cohort of cancer-diagnosed patients that NGS-based liquid biopsy can be used as a non-invasive adjunct tool for cancer monitoring in dogs. Surgically managed patients with no clinical residual disease but with a CSD result at the post-operative visit had a significantly higher probability of clinical recurrence within 6 months. By comparison, there was no significant difference in cancer recurrence within 6 months based on histologic margin assessment (complete vs. incomplete) alone. These findings suggest that pairing NGS-based liquid biopsy with surgical margin evaluation at the post-operative visit may provide a better indication of disease recurrence in the patient, and allow the clinician to develop a more personalized monitoring plan.

In surgically-managed patients that were followed longitudinally with liquid biopsy and had clinical recurrence of disease while undergoing monitoring, liquid biopsy detected molecular recurrence prior to clinical recurrence in 55% of patients (6/11; 95% CI: 25 – 82%), with a average lead time of over 4 months. This demonstrates that adding liquid biopsy to standard monitoring methods may aid in the early detection of cancer recurrence in some patients. Of note, the predefined time interval (i.e., 1 month after surgery or starting treatment and then every 3 months) for the follow-up visits (where clinical assessments and blood sample collection for liquid biopsy testing were performed) may have affected lead time estimates. The date of clinical recurrence for each patient was established based on the clinical disease status designation at these predefined visits and the medical records provided by the study sites. Some patients may have had a change in disease status at visits outside of the predefined visit schedule, so the actual date of clinical recurrence may have been different from the recorded date used in this analysis. More frequent liquid biopsy testing and clinical assessments may provide more accurate estimates for the lead time (from molecular recurrence to clinical recurrence) across various cancer types and therapies.

The goals of early identification of residual disease or disease recurrence during cancer monitoring are to achieve better disease control and improve patient outcomes by taking an individualized approach to patient care. In human medicine, results of liquid biopsy are currently being used for post-treatment cancer monitoring and to counsel patients about the risk of cancer recurrence.^15^ In some centers, liquid biopsy is also being used to personalize the patient’s treatment plan^16^ – tailoring therapies based on the genomic profile of the patient’s cancer, with the potential to move from a model of reactive treatment (i.e., initiating or changing treatment once clinical disease is evident) to that of proactive intervention (i.e., initiating or changing treatment when molecular evidence of disease is identified, even in the absence of clinical recurrence). Many clinical trials are ongoing to prospectively examine the effects of proactive intervention based on liquid biopsy results during cancer monitoring,^17^ and one trial recently showed that “ctDNA-positive patients appeared to derive considerable benefit from adjuvant treatment” when liquid biopsy was used to guide the initiation of such therapy in stage II colon cancer patients.^18^

In dogs, the clinical benefits of proactive intervention based on liquid biopsy results in the monitoring setting are yet to be established; therefore, specific recommendations for changes to patient treatment cannot yet be made. However, closer clinical monitoring of patients that receive a positive liquid biopsy result in the post-surgery or post-treatment setting appears prudent. For these patients, it may be reasonable to restage following a positive result, focusing on the primary anatomic site of the patient’s cancer, as well as common metastatic sites associated with the patient’s cancer type. If re-staging has already been completed, options would include offering additional diagnostics and/or more frequent monitoring. As with any diagnostic tool, liquid biopsy faces certain limitations. In the cancer monitoring setting, liquid biopsy is intended to be used as an adjunct to, rather than a replacement for, standard-of-care monitoring procedures. False positive and false negative results can occur with the test, and it is important to note that the test cannot currently distinguish between recurrence of the patient’s original cancer and development of a new cancer in the patient. The test reports a CSD or CSND result and does not indicate the extent of disease or whether a particular therapy is effective in reducing tumor burden. In the post-diagnosis setting, liquid biopsy is most useful for the detection of residual disease and the detection of disease recurrence in patients who had a CSD result at baseline (prior to any therapeutic intervention). In the absence of a baseline sample, the relative reassurance of a negative result during cancer monitoring is directly related to the detection rate for that cancer type, as described in the CANDiD study.^12^ Lastly, significant trauma, including tissue damage secondary to surgical procedures, can result in the temporary release of high amounts of cfDNA from damaged or necrosed normal cells and/or tumor cells^19^; in dogs undergoing surgery for tumor removal, high release of normal (non-cancer) cfDNA from damaged healthy tissues immediately after the procedure could dilute the residual tumor cfDNA fraction and impair the test’s ability to detect a cancer signal. Previous studies in humans^20^ and in dogs^21^, have shown that cfDNA fragments are usually cleared from circulation within a few days. Therefore, it is recommended to wait a minimum of 3 days, but conservatively 7 days, after surgery to perform liquid biopsy for the purpose of residual disease detection.

Longitudinal testing using NGS-based liquid biopsy has the potential to improve the clinical monitoring of dogs with cancer by detecting residual and recurrent disease with a simple blood draw. Patients with no clinical evidence of disease but with molecular residual disease or molecular recurrence detected by liquid biopsy may benefit from closer clinical evaluation and monitoring. Future studies are needed to evaluate the impact of liquid biopsy testing on clinical decision-making and on clinical outcomes across various cancer types and therapeutic interventions.

## Acknowledgements

The authors thank all the dogs involved in this study and the humans who love and care for them. The authors also thank the doctors and staff of the clinics that participated in the CANDiD study for the collection of samples and data that made this study possible. Additionally, the authors thank Lauren Wenstad for assistance with developing the tables, figures, and graphics in this manuscript, as well as the entire clinical studies and laboratory teams at PetDx for assistance with data generation.

